# Bayesian phylodynamic inference with complex models

**DOI:** 10.1101/268052

**Authors:** Erik M. Volz, Igor Siveroni

## Abstract

Population genetic modeling can enhance Bayesian phylogenetic inference by providing a realistic prior on the distribution of branch lengths and times of common ancestry.The parameters of a population genetic model may also have intrinsic importance, and simultaneous estimation of a phylogeny and model parameters has enabled phylodynamic inference of population growth rates, reproduction numbers, and effective population size through time. Phylodynamic inference based on pathogen genetic sequence data has emerged as useful supplement to epidemic surveillance, however commonly-used mechanistic models that are typically fitted to non-genetic surveillance data are rarely fitted to pathogen genetic data due to a dearth of software tools, and the theory required to conduct such inference has been developed only recently. We present a framework for coalescent-based phylogenetic and phylodynamic inference which enables highly-flexible modeling of demographic and epidemiological processes. This approach builds upon previous structured coalescent approaches and includes enhancements for computational speed, accuracy, and stability. A flexible markup language is described for translating parametric demographic or epidemiological models into a structured coalescent model enabling simultaneous estimation of demographic or epidemiological parameters and time-scaled phylogenies. We demonstrate the utility of these approaches by fitting compartmental epidemiological models to Ebola virus and Influenza A virus sequence data, demonstrating how important features of these epidemics, such as the reproduction number and epidemic curves, can be gleaned from genetic data. These approaches are provided as an open-source package *PhyDyn* for the BEAST phylogenetics platform.

## Introduction

Mechanistic models guided by expert knowledge can form an efficient prior on epidemic history when conducting phylodynamic inference with genetic data [1]. Parameters estimated by fitting mechanistic models, such as the reproduction number *R*_0_, are important for epidemic surveillance and forecasting. Compartmental models defined in terms of ordinary or stochastic differential equations are the most common type of mathematical infectious disease model, but in the area of phylodynamic inference, non-parametric approaches based on skyline coalescent models [2] or sampling-birth-death models [3] are more commonly used. Methods to translate compartmental infectious disease models into a population genetic framework have been developed only recently [4–8]. We address the gap in software tools for epidemic modeling and phylogenetic inference by developing a BEAST package, *PhyDyn*, which includes a highly-flexible mark-up language for defining compartmental infectious disease models in terms of ordinary differential equations. This flexible framework enables phylodynamic inference with the majority of published compartmental models, such as the common susceptible-infected-removed (SIR) model [9] and its variants, which are often fitted to non-genetic surveillance data. The *PhyDyn* model definition framework supports common mathematical functions, conditional logic, vectorized parameters and the definition of complex functions of time and/or state of the system. The *PhyDyn* package can make use of categorical metadata associated with each sampled sequences, such as location of sampling, demographic attributes of an infected patient (age, sex), or clinical biomarkers. Phylogeographic models designed to estimate migration rates between spatial demes [10–12] are special cases within this modeling framework, and more complex phylogeographic models (e.g. time-varying or state-dependent population size or migration rates) can also be easily defined in this framework.

The development of *PhyDyn* was influenced by and builds upon previous efforts to incorporate mechanistic infectious disease models in BEAST. The *bdsir* BEAST package [13] implements a simple SIR model which is fitted using an approximation to the sampling-birth-death process. The *phylodynamics* BEAST package [14] includes simple deterministic and stochastic SIR models which can be fitted using coalescent processes. More recently, the *EpiInf* package has been developed which can fit stochastic SIR models using an exact likelihood with particle filtering [15]. These epidemic modeling packages are, however, limited to unstructured populations (no spatial, risk-group, or demographic population heterogeneity). Other packages have been developed for spatially structured populations with a focus on phylogeographic inference, especially with the aim of estimating pathogen migration rates between discrete spatial locations [16]. The *MultiTypeTree* BEAST package [10] implements the exact structured coalescent model with multiple demes and with constant effective population size in each deme and constant migration rates between demes. Two BEAST packages, *BASTA* [17] and *MASCOT* [11] have been independently developed to use fast approximate structured coalescent models.These packages mirror the functionality of *MultiTypeTree* but include approximations to reduce computational requirements, enabling estimation of time-invariant effective population sizes and migration rates between spatial demes.

The *PhyDyn* BEAST package provides new functionality to the BEAST phylogenetics platform by implementing a much more complex family of structured coalescent models. In a general compartmental model, neither the effective population size nor migration rate between demes need be constant, and in more general frameworks, coalescence is also allowed between lineages occupying different demes. The package includes a flexible mark-up language for compartmental models including common mathematical functions making it simple to develop models which incorporate seasonality or which deviate from the simplistic mass-action premise of basic SIR models. The *PhyDyn* model mark-up language supports vectorised parameters (e.g. an array of transmission rates or population sizes) and simple conditional logic statements, so that epidemic dynamics can change in a discrete fashion, such as from year to year or in response to a public-health intervention. Commonly used phylogeographic models based on the structured coalescent are a special case of the general compartmental models implemented in the *PhyDyn* package, and extensions to the basic phylogeographic model can be implemented, such as by allowing effective population size to vary through time in each deme according to a mechanistic model.

## Design and Implementation

In this framework, first described in [5], we define deterministic demographic or epidemiological processes of a general form which includes the majority of compartmental models used in mathematical epidemiology and ecology. Defining compartmental models within this form facilitates interpretation of the population genetic model developed in the next section. Let there be *m* demes, and the population size within each deme is given by the vector-valued function of time *Y*_1:*m*_(*t*). We may also have *m*′ dynamic variables 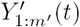 which are not demes (hence do not correspond to the state of a lineage), but which may influence the dynamics of *Y*. The dynamics of *Y* arise from a combination of *births* between and within demes, *migrations* between demes, and *deaths* within demes. We denote these as deterministic matrix-valued functions of time and the state of the system, following the framework in [5]:

- Births: *F*_1:*m*,1*:m*_(*t,Y,Y’*). This may also correspond to transmission rates between different types of hosts in epidemiological models.
- Migrations: *G*_1:*m*,1*:m*_(*t,Y,Y’*). These rates may have non-geographic interpretations in some models (e.g. aging, disease progression).
- Deaths: *µ*_1:*m*_(*t*, *Y*,*Y’*). These terms may also correspond to recovery in epidemiological models.

The elements *F*_*kl*_(…) describe the rate that new individuals in deme *l* are generated by individuals in deme *k.* For example, this may represent the rate that infected hosts of type *k* transmit to susceptible hosts of type *l*. The elements *G*_*kl*_(…) represent the rate that individuals in deme *k* change state to type *l*, but these rates do not describe the generation of new individuals. With the above functions defined, the dynamics of *Y*(*t*) can be computed by solving a system of *m* + *m*’ ordinary differential equations:

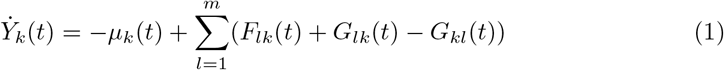

The *PhyDyn* package model markup language requires specifying the non-zero elements of *F*(*t*), *G*(*t*) and *µ*(*t*). There are multiple published examples of simple compartmental models developed in this framework [18-23]. In the following sections, we give examples of simple compartmental models related to infectious diease dynamics and show how these models can be defined within this framework. We provide examples of models fitted to data from seasonal human Influenza virus and Ebola virus as well as a simulation study.

### Seasonal human Influenza model

We model a single season of Influenza A virus (IAV) H3N2 and apply this model to 102 HA-1 sequences collected between 2004 and 2005 in New York state [24, 25]. We build on a simple susceptible-infected-recovered (SIR) model which accounts for importations of lineages from the global reservoir of IAV, which we will see is a requirement for good model fit to these data (Figure 1). This model has two demes: The first deme corresponds IAV lineages circulating in New York, and the second deme corresponds to the global IAV reservoir. The global reservoir will be modeled as a constant-size coalescent process. Within New York state, new infections are generated at the rate *βI*(*t*)*S*(*t*)/*N* where *β* is the per-capita transmission rate, *I*(*t*) is the number of infected and infectious hosts, *S*(*t*) is the number of hosts susceptible to infection, and *N = S + I + R* is the population size. *R*(*t*) denotes the number of hosts that have been infected and are now immune to this particular seasonal variant. With the above definitions, we define the matrix-valued function of time:

**Fig 1.**
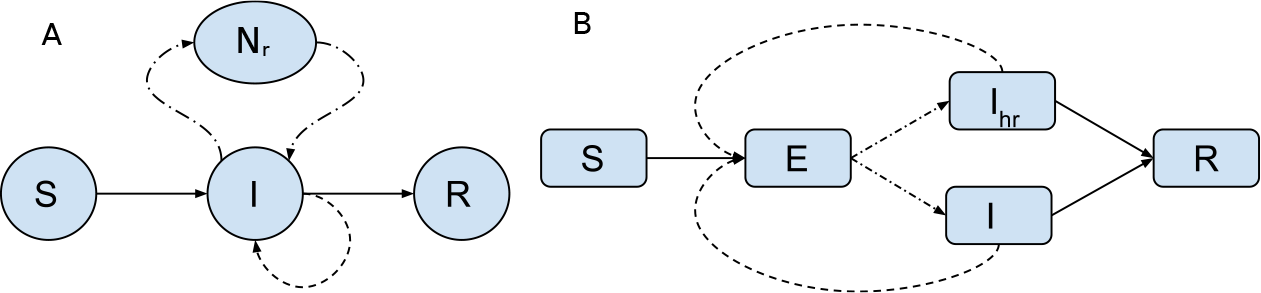
Compartmental diagram representing structure of models for seasonal human Influenza (A) and Ebola virus models (B and C). Solid lines represent flux of hosts between different categories. Dash lines represent migration. Dotted lines represent births (transmission).

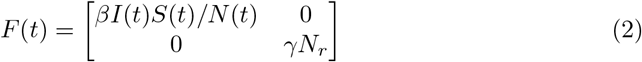

Note that births within the reservoir do not vary through time and depend on the effective population size in that deme *N*_*r*_.

Additionally, we model *deaths* from the pool of infected using

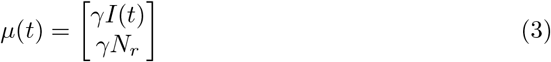

Births balance deaths in the reservoir population.

Finally, we model a symmetric migration process between the reservoir and New York:

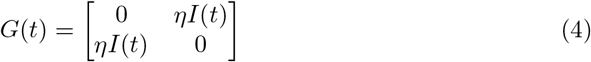

where *η* is the per-capita migration rate. Note that migration between the reservoir and New York are balanced and do not effect the dynamics of *I*(*t*) over time.

These three processes lead to the following differential equation for the dynamics of *I*(*t*)

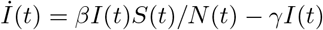

Below, we show a fit of this model where the following parameters are estimated:

- Migration rate *η;* prior: lognormal (log mean=1.38, log sd = 1)
- Recovery rate *γ*; prior: lognormal(log mean = 4.8, log sd = 0.25)
- Reproduction number *R*_0_ = *β/γ;* prior: lognormal (log mean 0, log sd = 1)
- Reservoir size *N*_*r*_; prior: lognormal (log mean = 9.2, log sd = 1)
- Initial number infected in September 2004; prior: lognormal (log mean = 0, log sd = 1)
- Initial number susceptible in September 2004; lognormal (log mean = 9.2, log sd =1)

Note that the model only had one informative prior, which was for the recovery rate, and was based on the previous study of viral shedding by Cori et al. [26]

### Ebola Virus in Western Africa

We develop a susceptible-exposed-infected-recovered (SEIR) model (Figure 1) for the 2014-2015 Ebola Virus (EBOV) epidemic in Western Africa and apply this model to phylogenies previously estimated by Dudas et al. [27]. Phylogenies estimated by Dudas are randomly downsampled to *n* = 400 to alleviate computational requirements.

According to the SEIR model, infected hosts progress from an uninfectious exposed state (E) to an infectious state (I) at rate *γ*_0_ which influences the generation-time distribution of the epidemic. Infectious hosts die or recover at the rate *γ*_1_. The SEIR model has the following form:

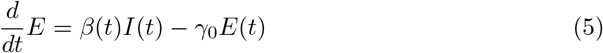

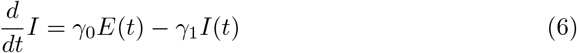

where *β*(*t*) is the per-capita transmission rate. In a typical mass-action model, we would have *β*(*t*) ∝ *S*(*t*)/(*S*(*t*) + *E*(*t*) + *I*(*t*) + *R*(*t*)), however in order to demonstrate the flexibility of this modeling framework, we will instead use a simple linear function, *β*(*t*) = *at* + *b,* and in general a wide variety of parametric and non-parametric functions could be used within the BEAST package to model the force of infection.

There are two demes in this model corresponding to the potential states of an infected hosts. The birth matrix with demes in the order (*E, I*) is

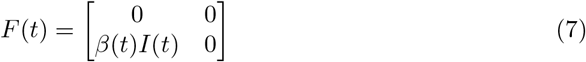

The migration matrix encapsulates all processes which may change the state of a lineage without leading to coalescence of lineages, and this includes progression from E to I:

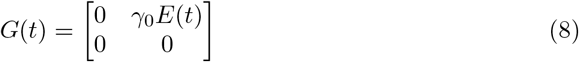

And finally removals are modeled using

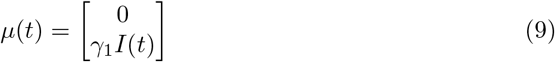

Note that the parametric description of *β*(*t*) does not require us to model dynamics of *S*(*t*) or *R*(*t*).

The parameters estimated and priors for this model are

- *β*(*t*) slope a, prior: Normal(0, 40)
- *β*(*t*) intercept b, prior: lognormal(log mean = 4.6, log sd = 1)
- Initial number infected (beginning of 2014), prior: lognormal (log mean=0, log sd = 1)

In order to reconstruct an epidemic trajectory which closely matched the absolute numbers of cases through time, we include additional variables that could influence the relationship between effective population size and the true number of infected hosts. For this purpose we developed a second EBOV model which included higher variance in the offspring distribution, reasoning that a higher variance in the number of transmissions per infected case would lead to higher estimates of the epidemic size [28]. The superspreading model (Figure 1) includes two infectious compartments, *I*_*l*_ and *I*_*h*_, with per-capita transmission rates *β*(*t*) and *τβ*(*t*) respectively. The factor of *τ* > 1 represents a transmission risk ratio for the second infectious deme. We specify that a constant fraction *p*_*hr*_ progress from *E* to *I*_*h*_, with the remainder going to *I*_*l*_. With demes in the order (*E, I*_*l*_, *I*_*h*_), the birth, migration, and death matrices for the superspreading model are as follows:

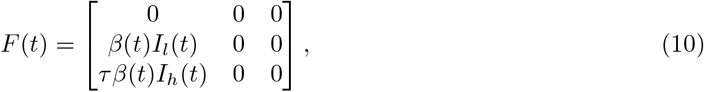

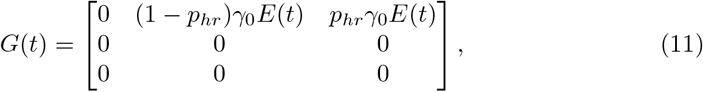

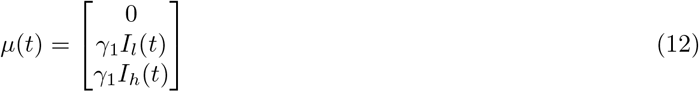

Additional parameters and priors for the superspreading model are

- *τ*, prior: lognormal (log mean = 1, log sd = 1)
- *p_hr_,* fixed at 20%

### Simulation model

We developed a simulation model with four demes in order to evaluate the ability of BEAST to identify and estimate birth rates, migration rates, and transmission risk ratios. This model includes two types of hosts, with low and high transmission risk. Additionally, each type of host progresses through two stages of infection, where the first stage is short but has higher transmission rate. The four demes are denoted *Y*_0*l*_, *Y*_1*l*_, *Y*_0_*h, Y*_1*h*_ where the first subscript denotes stage of infection and the second subscript denotes transmission risk level. The birth matrix is

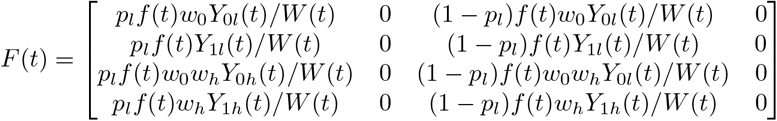

In this model, a proportion *p*_*l*_ of all transmissions go to the low risk group. Transmissions from stage 1 are proportional to the transmission risk ratio *w*_0_ > 1. Transmissions from the high risk group are proportional to the transmission risk ratio *w*_*h*_ > 1. The variable *W*(*t*) = *w*_0_*Y*_0*l*_ *+ Y*_1*l*_ *+ w*_0_*w*_*h*_*Y*_0*h*_ + *w*_*h*_*Y*_1*h*_ normalizes the proportion of transmissions attributable to each deme. The variable *f*(*t*) gives the total number of transmissions per unit time, and for this we use a SIRS model:

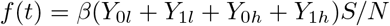

where *S*(*t*) is the number susceptible governed by the following equation

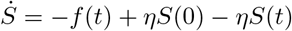

and *η* is the per-capita rate of non-disease related mortality.

The migration matrix captures the disease stage-progression process:

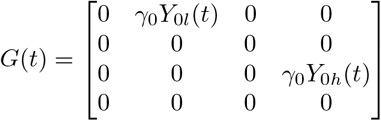

The death matrix is

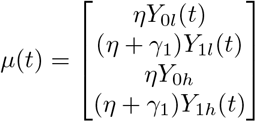

To generate simulated data, we simulated epidemics using Gillespie’s exact algorithm over a discrete population and an initial susceptible population of two thousand individuals. A random sample of *n* = 250 was collected between times 100 and 250 and the history of transmissions was used to reconstruct a genealogy. BEAST PhyDyn was then used to estimate

- *β,* prior: lognormal (log mean=-1.6, log sd = 0.5)
- *w*_0_, prior: uniform (0, 50)
- *w*_*h*_, prior: uniform (0, 50)
- The initial number infected, prior: lognormal (log mean=0, log sd = 1)

Note that BEAST PhyDyn is fitting deterministic models to data generated from a noisy stochastic process and some error should be expected due to this approximation. Supporting figure shows a comparison of a single noisy simulated trajectory and a solution of the deterministic model under the true parameters. All simulation code and BEAST XML files are available at https://github.com/emvolz/PhyDyn-simulations.

### Modeling the coalescent process conditioning on a complex demographic history

In this section we review the approximate structured coalescent model described in [5] and describe extensions designed to improve accuracy and reduce computational cost. A complete description of the structured coalescent, likelihood calculations, and approximations made in this framework are contained in the Supporting Information.

The probability that a lineage *i* in a bifurcating rooted genealogy 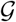 is in deme *k* ∈ 1 : m at time *t* will be denoted *p*_*ik*_(*t*). Usually the state of a lineage will be observed at the time of sampling *t*_*i*_, so that *p*_*ik*_(*t*_*i*_) is a point density. We compute the likelihood by solving a system of differential equations for the ancestral states *p*_1:(2*n*-2),1:*m*_ and computing the expected coalescent rate between each pair of lineages.

Let 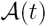 denote the set of extant lineages at time *t*. The expected number of lineages in each deme as a function of time is 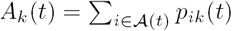.

Given lineages *i* and *j* ∈ 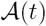, we model the rate of coalescence between the pair of lineages as a function of the ancestral state vectors *p*_*i*,1:*m*_(*t*) [5, 29]. Justification for this modeling approach is provied in the supporting information.

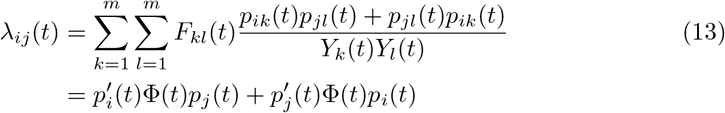

where Φ(*t*) is a *m* × *m* matrix with elements

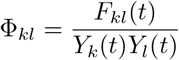

This matrix describes the rate that a lineage in deme *k* coalesces with a lineage in deme *l* as a result of birth events from the former to the latter. Note that *λ*_*ij*_ = *λ*_*ji*_. The total rate of coalescence at time *t* is

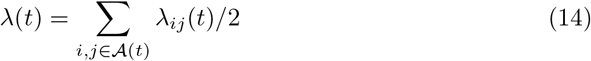

The probability 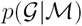 of a given labeled genealogy given a demographic history 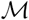 = (*F, G, Y*), described in previous publications [5, 29], is that of a point process with intensity *λ*(*t*) multiplied by the the multinomial density with probabilities *λ*_*ij*_(*t*)/*λ*(*t*) for all pairs of lineages *i* and *j* which coalesce. The form of this likelihood is further reviewed in the supporting information. Approximations to the structured coalescent differ in how ancestral state vectors are derived. In [5] in 2012, the following approximation (denoted “com12”) was presented which required the solution of the following differential equations:

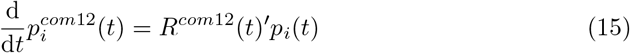

where *R* is the *m* × *m* matrix with elements

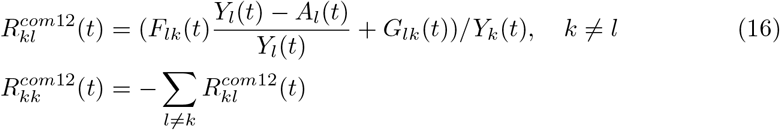

Note the inclusion of the term 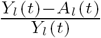 in equation 16, which was an approximation intended to account for the fact that only lineages not ancestral to the sample could cause a lineage to change state without resulting in coalescence. The form of this equation was found to provide an accurate approximation to the lineages through time in [5], when solving the system of equations

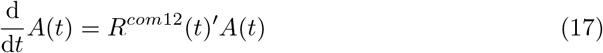

A drawback of the *com12* model is that it does not condition on all possible events that lineage *i* may experience during its evolution; namely it does not condition on the observation that no coalescent events occurred during an internode interval. Consequently, this model tends to overestimate the probability that a lineage occupies a deme with a high coalescent rate. This issue was thoroughly explored in a recent investigation by Muller et al. [30] in the special case of phylogeographic models (diagonal *F*(*t*), constant rates, constant population size). A preliminary model which addresses this issue in was also introduced by Volz in the *rcolgem* R package [31], and we build on the approach developed therein which explicitly computes the cumulative probability that lineage *i* has coalesced (see Supporting Information). To make this model tractable, we make the approximation that the marginal probabilities of each pair of lineages are independent. In other words, we make the approximation that the probability that lineage *i* is in deme *k* is independent of the probability that a different extant lineage *j* is in deme *l*. Let 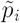 represent an augmented state vector with *m* + 1 elements where 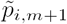 represents the probability that lineage *i* has coalesced. Note that *m* + 1 is an absorbing state which increases at the rate *λ*_*i*_. (*t*). In the Supporting Information, we show:

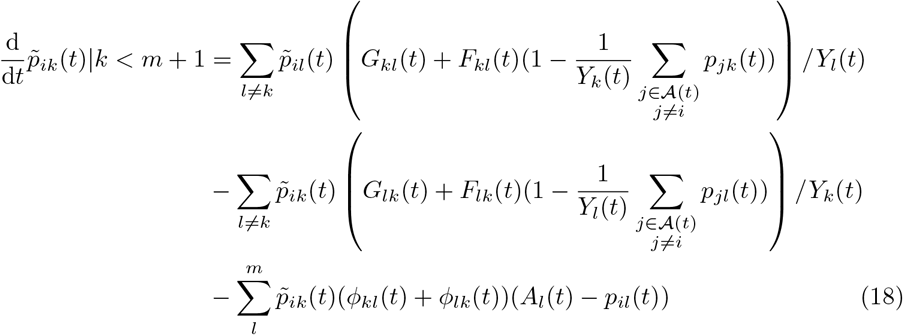

The first line represents a change in deme from *l* to *k* (due to birth or migration), the second line represents change from deme *k,* and the last line accounts for coalescence of lineage *i.*

Note that if the rate of coalescence is non-zero over the history a lineage, 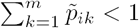. If the ancestor node of a lineage *i* occurs at a time *τ*, we derive *p*_*i*_(*τ*) by renormalizing the distribution computed from equation 18, which provides the state vector conditional on the event that no coalescence was observed:

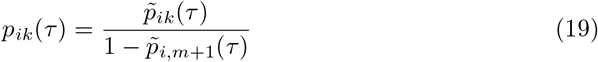

Unfortunately, the system of equations 18 can be slow to solve since it requires recursion over extant lineages (twice) and *m* + 1 ancestral states. We therefore provide an additional approximation which greatly reduces computational cost and is closely related to the approach described in [5]. We define *Q*(*t, T*) to be the *m × m* matrix of transition probabilities such that

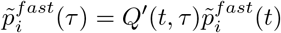

and 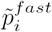 provides the length *m* unormalized state vector analaogous to 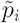 above. his vector can be renormalized so that 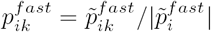. We can also approximate the number of lineages in each deme over the interval using

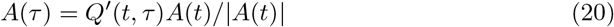

where *t* < *τ* < *T.* Finally, we can modify equation 18 to use the vector *A*(*τ*), avoiding the need to sum over extant lineages. The following matrix provides transition rates between states.

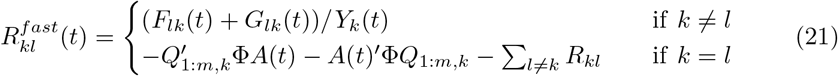

Note that the diagonal of the rate matrix also includes the approximate rate that a lineage will coalesce given that it begins the internode interval in each of the *m* demes. Thus this is a ‘leaky’ Markov process and the probabilities in each column of *Q* will not in general sum to one.

Initally *Q*(*t, t*) = *I* is the identity matrix. The matrix *Q*(*t*, *T*) is computed by solving the equations

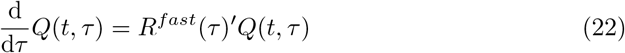

Therefore only *m*^2^ differential equations need to be solved over every internode interval. Note that the rate of coalescence in equation 21 exceeds that in 18, so that this fast approximation will slightly over-estimate the probability that a lineage coalesces over an interval.

We will denote the model based on equations 18 *comP* and the model based on equations 21 *comQ*. Performance of *comQ* and *comP* is explored in simulations below.

## Results

With simulated data, BEAST PhyDyn recovers the correct transmission risk ratios and transmission rates using both the *comP* model (equation 18) and the faster *comQ* model (21). Figure 2 compares estimates across twenty simulations using both variants. The running time of the *comQ* model was approximately five times faster than *comP* in these simulations with trees that have 250 samples and four demes. Good coverage of parameter estimates was observed with the *comP* model. Across 60 parameter estimates (three parameters not counting initial conditions and twenty simulations), estimates did not cover the true value two times. The *comQ* model failed to cover in five of 60 estimates. Greater bias was observed with the *comQ* model, with the greatest bias observed for the *w*_*h*_ parameter (cf equation 13, mean estimate 3.63, truth:5). However the *comQ* model also had superior precision, with a posterior root mean square error of 2.4 versus 4.8 observed with the *comP* model. A similar but less pronounced pattern of bias and precision was observed for other parameters. A complete summary of simulation results is available at https://github.com/emvolz/PhyDyn-simulations.

### Human Influenza A/H3N2

The seasonal influenza SIR model which accounts for importations from the global reservoir was applied to 102 HA/H3N2 sequences collected from New York state during the 2004-2005 flu season. These data were previously analyzed using non-parametric models by [24]. Figure 3 shows the estimated posterior effective number of infections over the course of the influenza season, and the time of peak prevalence is correctly identified around the end of 2004. We also compared the model-based estimates to non-parametric estimates generated in BEAST using a conventional non-parametric Bayesian skyline model which is also shown in Figure 3. The skyline model does not detect a decrease in prevalence towards the end of the influenza season and does not identify the time of peak prevalence. Use of a well-specified parametric compartmental model imposes a strong prior on the epidemic trajectory which leads to the correct identification of the shape and timing of the epidemic curve.

We estimated the reproduction number *R*_0_ = 1.16 (95%CI: 1.07-1.30). This value is similar to many previous estimates based on non-genetic data for seasonal influenza in humans which according to the recent review in [32] have an interquartile range of 1.18-1.27 for H3N2. Bettancourt et al. [33] estimated *R*_0_ = 1.22 for the 2004-05 H3N2 seasonal influenza epidemic in the entire USA using weekly case report data. An uninformative prior was used for *R*_0_ in the BEAST/PhyDyn analysis.

### Ebola virus in Western Africa

We applied the SEIR and superspreading-SEIR models to Ebola virus phylogenies based on data first described by [27] and subsequently analyzed in [34]. These phylogenies were estimated from whole genome sequences collected 2014-2015 during the West African Ebola epidemic. We derived the maximum clade credibility tree from the analysis by [27] and extracted a subtree based on sampling four hundred lineages at random. The BEAST PhyDyn package was used to fit the models with fixed tree topologies and branch lengths. We also ran the analysis using a fixed tree estimated by maximum likelihood and the *treedater* R package as described in [34], finding similar results.

**Fig 2.**
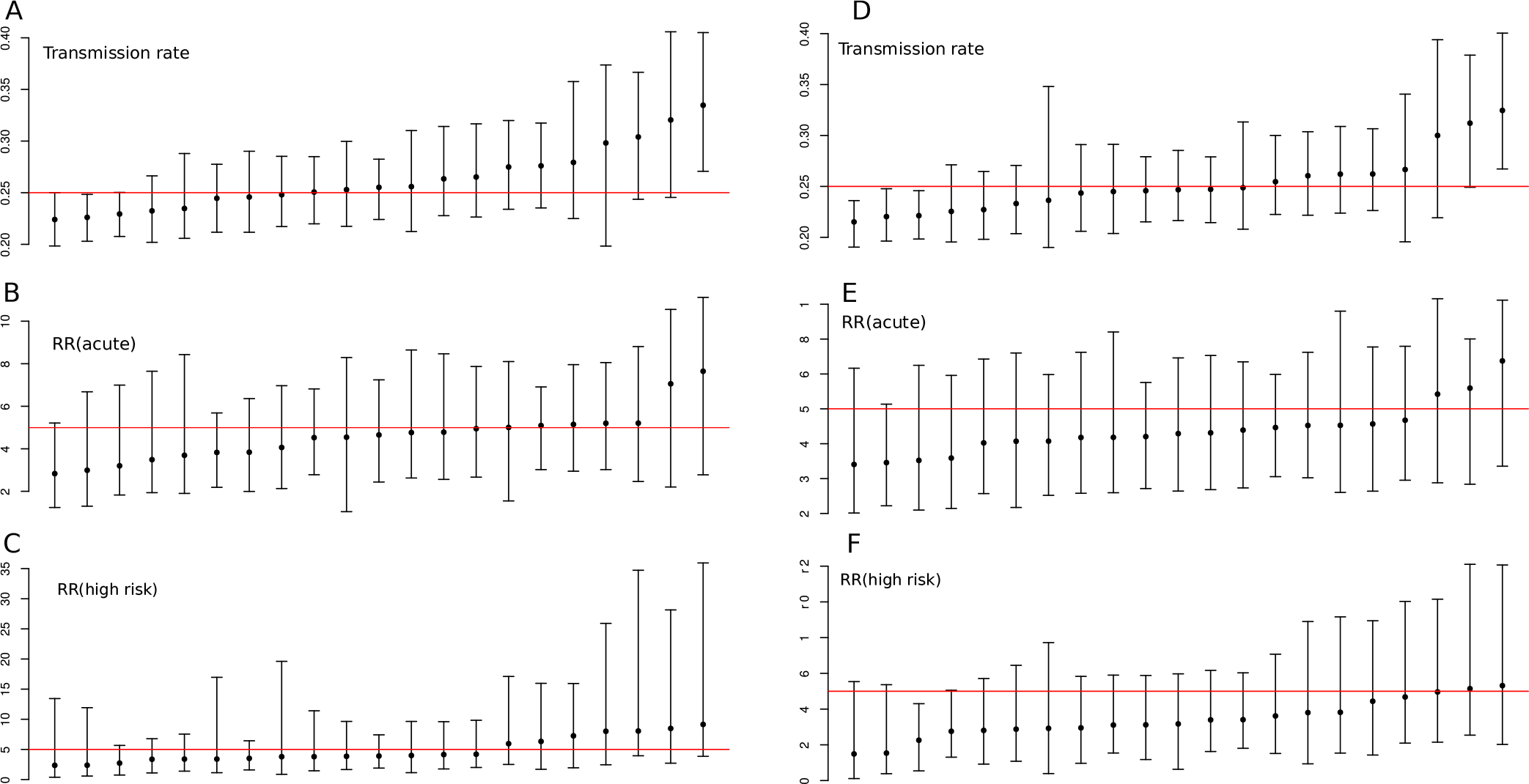
Parameter estimates and credible intervals for 20 simulations. The red line shows the true value. A-C: Results generated using the *comP* model. D-F: Results generated using the *comQ* model.

**Fig 3.**
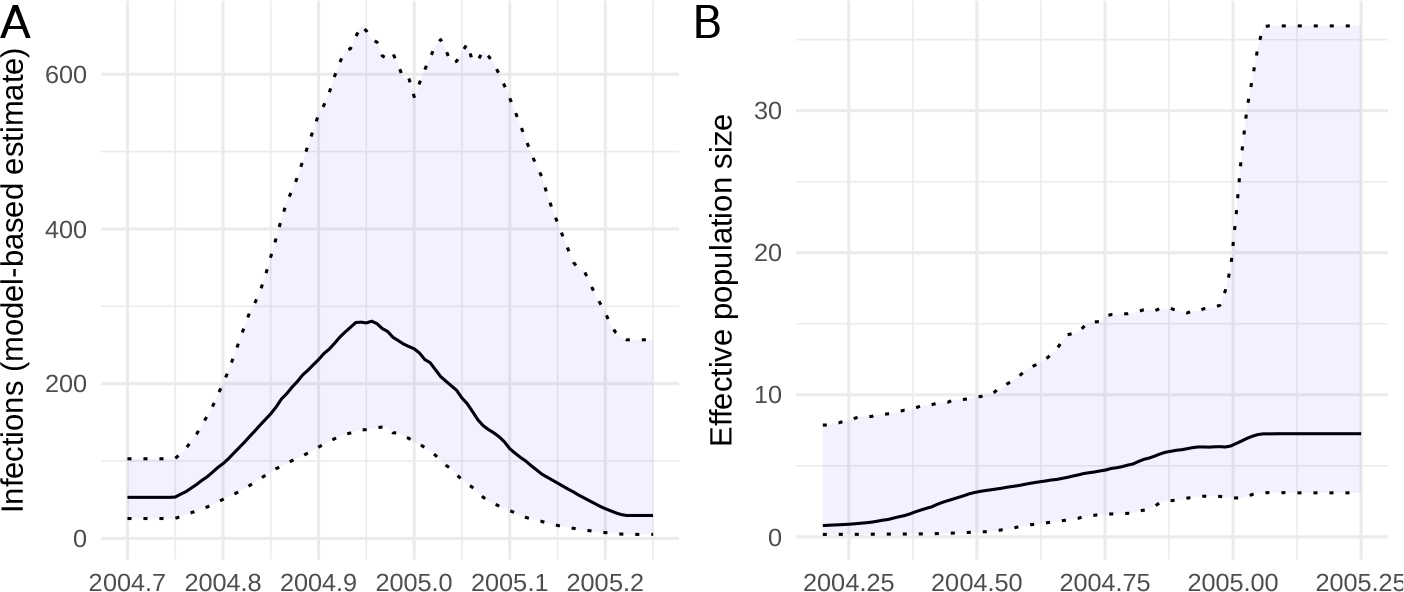
The estimated effective number of H3N2 human influenza infections in 2004-2005 in New York State. A. Estimates obtained using the parametric seasonal influenza model described in the text. B. Effective population size estimated using a conventional Bayesian skyline analysis.

**Fig 4.**
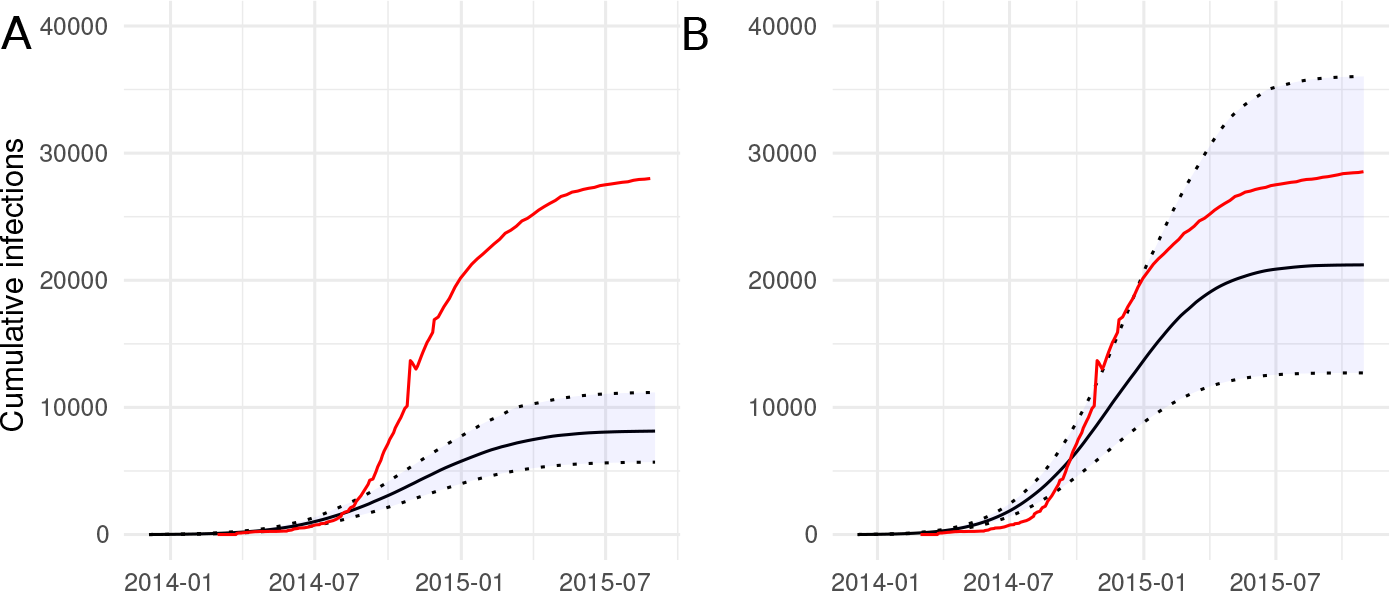
Model-based estimates of cumulative infections through time for the 2014–15 Ebola epidemic in Western Africa. Estimates are shown for the SEIR model (A) and the model which includes super-spreading (B). The red line show the cumulative number of cases reported by WHO [34].

**Fig 5.**
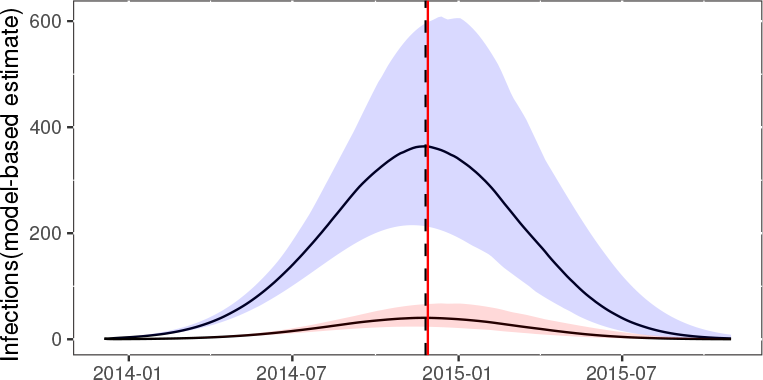
Estimated effective number of infections through time using the superspreading SEIR model for the 2014-15 Ebola epidemic in Western Africa. The red vertical line shows the time of peak prevalence inferred from WHO case reports. The vertical dashed line shows the model estimated time of peak prevalence. The red trajectory shows the proportion of infections in the high-transmission-rate compartment.

We estimated similar reproduction numbers using both models. With the SEIR model, we estimate *R*_0_ = 1.47(95%CI: 1.41-1.53). With the superspreading-SEIR model, we estimate *R*_0_ = 1.52(95%CI:1.48-1.54). Note that uninformative priors were used for parameters determining *R*_0_. As anticipated, the model fits provide substantially different estimates of the cumulative number of infections. Figure 4 shows the estimated cumulative infections through time using both models alongside the cumulative number of cases reported by WHO and compiled by the US CDC [34]. Both models provide similar estimates regarding the relative numbers infected through time and the time of epidemic peak. Using the superspreading model, the time of peak incidence is estimated to have occurred on November 25, 2014. According to WHO reports, this occurred only three days later on November 28 (Figure 5.

Estimates of cumulative infections with the superspreading model are consistent with WHO data, whereas results with the SEIR model are not. The superspreading model accomodates an over-dispersed offspring distribution (the number of transmission per infection), thereby decreasing effective population size per number infected and yielding larger estimates for the number infected [28]. We estimate the transmission risk ratio parameter (ratio of transmission rates between high and low compartments) to be 8.1 (95%CI: 6.68-10.73). This implies that a minority of 10% of infected individuals are responsible for 43%-54% of infections.

Formal model comparison methods such as Bayesian stepping-stone approaches [35] are not yet supported by the *PhyDyn* package, but we note that a much higher mean posterior likelihood was found using the superspreading model (−1006.9) than with the SEIR model (−1068.5).

## Availability and Future Directions

The *PhyDyn* package, source code, documentation and examples can be found at https://github.com/mrc-ide/PhyDyn. The *PhyDyn* package greatly expands the range of epidemiological, ecological, and phylogeographic models that can be fitted within the BEAST Bayesian phylogenetics framework. Extensions enabled by this package include models with parametric seasonal forcing, non-constant parametric migration or coalescent rates between demes, state-dependent migration or coalescent rates, and discrete changes in migration or coalescent rates in response to perturbation of the system (e.g. a public health intervention). The package also provides a means of utilizing non-geographic categorical metadata which is usually not considered in phylodynamic analyses, such as clinical or demographic attributes of patients in a viral phylodynamics application [19].

We have demonstrated the utility of this framework using data from Influenza and Ebola virus epidemics in humans, finding epidemic parameters and epidemic trajectories consistent with other surveillance data. In both of these examples, simple structured models were fitted, but notably without using any categorical metadata associated with sampled sequences. This demonstrates potential advantages of structured coalescent modeling even in the absence of informative metadata. In the case of human Influenza A virus, the fitted model included a deme which accounted for evolution in the unsampled global influenza reservoir, which allowed estimation of epidemic parameters within the smaller sub-region which was intensively sampled. The use of a parametric mass-action model allowed *PhyDyn* to correctly detect the time of epidemic peak and epidemic decline, whereas non-parametric skyline methods did not detect epidemic decline in this case. And in the application to the Ebola virus epidemic in Western Africa, models included un-sampled ‘exposed’ categories which accounted for realistic progression of disease among patients, as well as a ‘super-spreading’ compartment which accounted for over-dispersion in the number of transmissions per infected case.

In developing *PhyDyn*, the focus has been on developing a highly flexible framework which is also computationally tractable for moderate sample sizes and model complexity. But flexibility and computational efficiency has come at the cost of some realism, notably in the deterministic nature of the models included in this framework. Future extensions may utilize stochastic epidemic models such as those described by [29]. Other directions for future development include semi-parametric modeling, such as models with a spline-valued force of infection [22] or models utilizing Gaussian processes [36], and approaches for utilizing continuous-valued metadata [37].

## Supporting information

### Structured coalescent likelihood and approximations

The structured coalescent defines a discrete Markovian stochastic tree-building process in retrospective/continuous time [38]. In it’s most general form, the process defines the potentially time and state dependent rates that 1) a lineage changes demes and 2) the rate of ‘coalescence’, i.e. the rate that two or more lineages form a single lineage at a node. In this section, we explain how the (*F, G, Y*) model-based coalescent is related to the more general process and approximations involved in computing the fast likelihood. We consider a structured coalescent for a bifurcating genealogy that has the following events and rates:

- Deme change. A lineage changes deme from *k* to *l* at the rate *M*_*kl*_(*t*).
- Coalescence. A lineage in deme *k* ‘gives birth’ to a lineage in deme l at the rate *φ*_*kl*_(*t*) giving rise to an ancestor lineage in deme *k.*

The vast majority of previous work on these models has been focused on phylogeographic applications and therefore consider only the special case *ϕ*_*kl*_ = 0 if *k* ≠ *l,* i.e. lineages only coalesce if they are co-located in the same deme. In these models, we depart from the usual approach and specify different coalescence rates ϕ*kl* and *ϕ*_*lk*_ which yield different types of ancestor lineages if *k ≠ l.* Note that the total rate of coalescence of a lineage *i* in deme *k* and a lineage *j* in deme l would be *ϕ*_*kl*_ + *ϕ*_*lk*_. Most previous work has also focused on the special case of constant rates, so that *M* and *ϕ* are independent of time. Epidemiological and ecological models will often have both time-dependent rates and non-zero off-diagonal elements in *ϕ*.

At any time in the course of the tree-building process, a set of ‘extant’ lineages 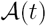 with cardinality *A*(*t*) will have configuration *S*(*t*) = (*s*_1_,…, *s*_*A*(*t*))_ where *s*_*i*_ ∈ 1 : *m* is an integer index of *m* demes. We will also use the notation *A*_*k*_(*t*) to denote the number of extant lineages in deme *k*. The likelihood of a genealogy under the structured coalescent is defined as that of a point process with competing rates for different types of events (coalescence or deme-change). Note that we will also allow rates of coalescence and deme-change to be time and state-dependent. The probability density of a genealogy produced by this process is easily computed provided that the configuration *S*(*t*) is known at all time points [12, 15].

Additional notation is required to specify this density:

- In a rooted bifurfacting genealogy with *n* tips, there are 2*n* − 2 lineages. We index these such that *i* ∈ 1: *n* corresponds to terminal branches of the genealogy, and *i* ∈ (*n* + 1) : (2*n* − 2) is ancestral to an internal node of the tree.
- 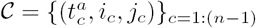 is the set of internal nodes represented by a tuple and each also corresond to coalescent events. The indices *i* and *j* correspond to the daughter lineages of the internal node, and *t*^*a*^ is the time when the internal node occurs
- *T*_*S*_ = (*t*_1_, …, *t*_*n*_) denotes times of sampling. We assume the state at time of sampling *s*_*i*_(*t*_*i*_) is known.
- 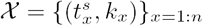 denotes the set of sample events represented by a tuple. The time of sampling is *t*^*s*^ and the observed deme at time of sampling is *k.*
- 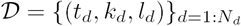 denotes the set of *N*_*d*_ deme-change events represented by a tuple. The time of the event is denoted *t*_*d*_. The initial deme is *k* and the deme following the change is *l*.
- Let *T =* (t_1_, …, *t*_*N*_) denotes the ordered sequence of all event times (sampling, migration, or coalescence) in the tree building process.

Note that the total coalescence rate between lineages *i* and *j* given known lineage-deme configuration *S* is

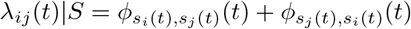

We can define the cumulative hazard of migration or coalescence between consecutive time points in *T.* Note that the configuration *S* is fixed during this period, but migration and coalescence rates may vary. The cumulative hazard over an interval is:

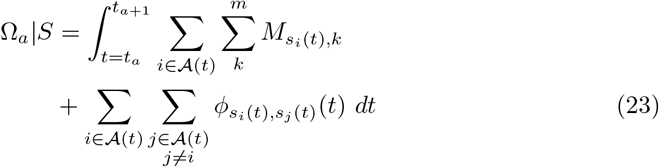

The joint probability density of the genealogy and deme-configuration *S*(*t*) can now be defined:

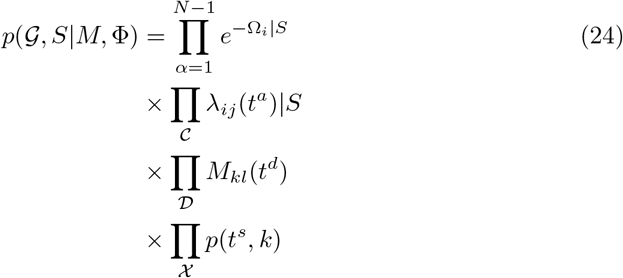

The first line provides the probability of no events occurring in each internode interval, the second and third lines provide the rates of each coalescent and deme-change event, and the last line describes the density of the sample times and sample states [39]; in most applications there is not a model of the sampling process or prior information to establish this density, and it is not included.

Since in practice the configurations *S*(*t*) are not observed except at times of sampling, estimation of migration rates and/or effective population sizes then requires a strategy for marginalizing over latent *S*(*t*), and this is typically done by MCMC simulation [15]. This entails a large computational cost of integrating over a vary large configuration space [17], so instead we pursue a strategy of developing equations to marginalize over lineage states and will derive an approximate likelihood in terms of the probability that a lineage is in a particular deme. Our strategy is to derive equations for *p*_*ik*_(*t*) = *p*(*s*_*i*_(*t*) = *k*) at all times *t* over the history of each lineage *i.* Deriving equations for the joint probability of *s*_*i*_ and *s*_*j*_ is complicated, so we make the simplifying approximation that the conditional probability of state *j* is equal to the marginal: *p*(*s*_*j*_ = *l*|*s*_*i*_ = *k*) *≈ p*(*s*_*j*_ = *l*). We may then compute rates of deme-change for lineage *i* and the rate of coalescence of *i* with all other lineages. Where the configuration is unknown, we have

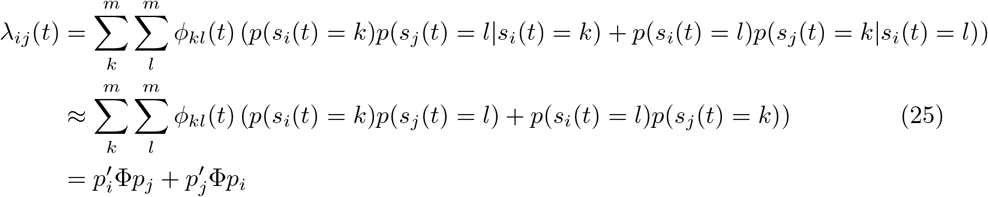

We compute the cumulative hazard of coalescence in internode intervals as before, but considering only times of sampling and times of coalescence. In contrast to the above equation for Ω_*i*_|*S*, the lineage configuration will change over the course of each internode interval. The symbol *S*(*t*) denotes the set of all possible lineage-deme configurations at time *t* and *p*(*S*(*t*)) is the marginal probability of a particular configuration *S*.

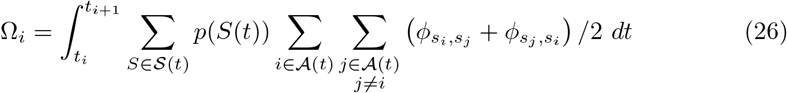

We introduce the indicator function *I*(*x* = *y*) which evaluates as one if and only if *x* equals *y* and is zero otherwise. Rearranging the order of summation

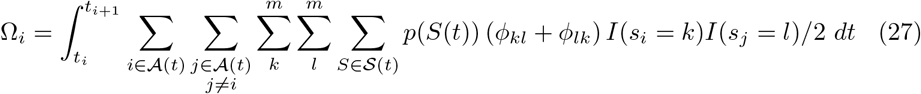

Recognizing that 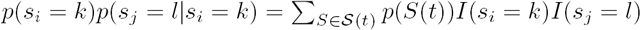, we can simplify the preceding:

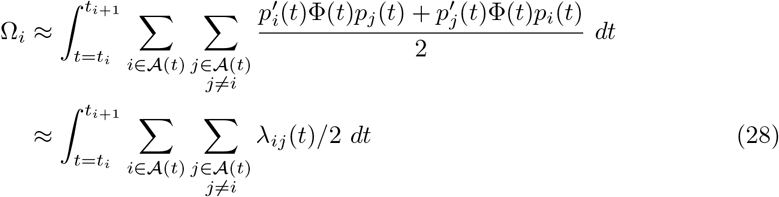

Thus we can avoid computing the full intractable density *p*(*S*(*t*)) and use the approximation for *λ*_*ij*_(*t*) in equation 25 provided that we can solve for the marginal probabilities *p*(*s*_*i*_ = *k*) and it is shown how to do that below.

Marginalising over all possible configurations in *S*(*t*), the density of a genealogy is

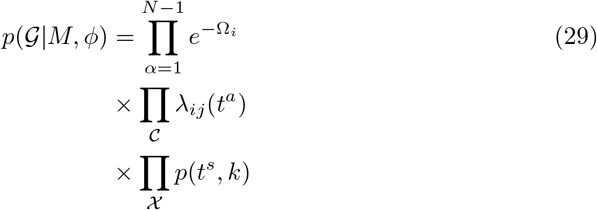

We now derive a system of *m* + 1 equations for the probability *p*_*ik*_(*t*) = *p*(*s*_*i*_(*t*) = *k*). These equations are defined in terms of the (*F, G, Y*) model decomposition and the reader should review the main text and reference [5] for how this works. As stated here, these equations are self-conistent, and it is only of importance that the rates of different migration and coalescent events through time are pre-specified. The events which can modify the deme of lineage *i* when tracing its history backwards in time are

- Migration. Lineage *i* will move from deme *k* to *l* at a rate *G*_*lk*_(*t*)/*Y*_*k*_(*t*).
- Birth. A lineage not ancestral to the sample (not represented by a lineage in the tree) may give birth to lineage *i.* The deme of lineage *i* would then be the deme of the lineage which gave birth to *i.*
- Coalescence. We must condition on the event that no coalescence is observed over each internode interval.

We find it necessary to distinguish between birth events which result in coalescence and birth events which will not. Note that this is not usually required in phylogeographic coalescent models because birth events would not result in deme change but only in potential coalescence. Collecting the first two types of events which change the deme of a lineage, we can define the following time dependent migration matrix for lineage *i:*

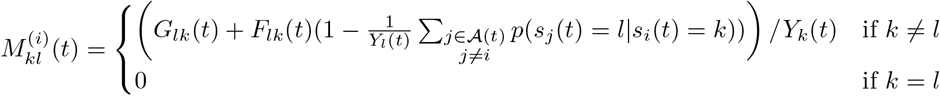

This provides the rate that lineage i migrates from deme *k* to *l* at time *t*. As above, the joint density for *s*_*i*_ and *s*_*j*_ is not tractable, so we use the approximation

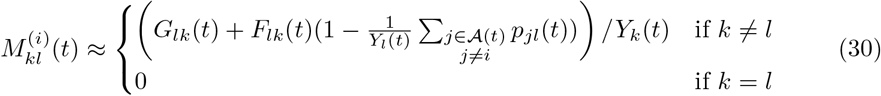

To account for the effect of coalescence on evolution of the state vector *p*_*i*_(*t*), we instead develop equations for the augmented state vector 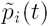, which has *m* + 1 elements and where 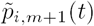 represents the probability that lineage *i* has coalesced at some time 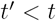. If *t*_*i*_ represents the initial time of lineage *i* (the most recent time that *i* exists on forward time axis), we have

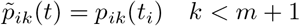

and 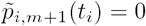. This vector provides the marginal probability of a ‘leaky’ continuous time Markov process with *m* + 1 states, where *m* + 1 is an absorbing state. To retrieve the state vector conditioning on no coalescent event having occured, we use the identity

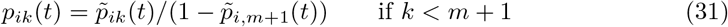

The forward equations for the evolution of 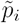 are found by tabulating the effects of migration, birth, and coalescent events outlined above:

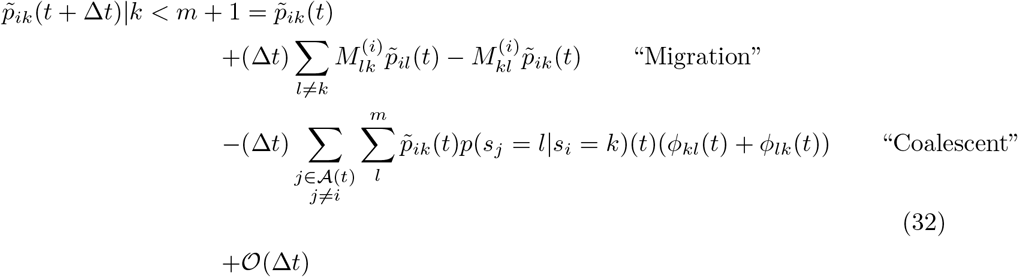

Using the approximation *p*(*s*_*j*_ = *l*|*s*_*i*_ = *k*) ≈ *p*_*jl*_(*t*) and simplifying this becomes

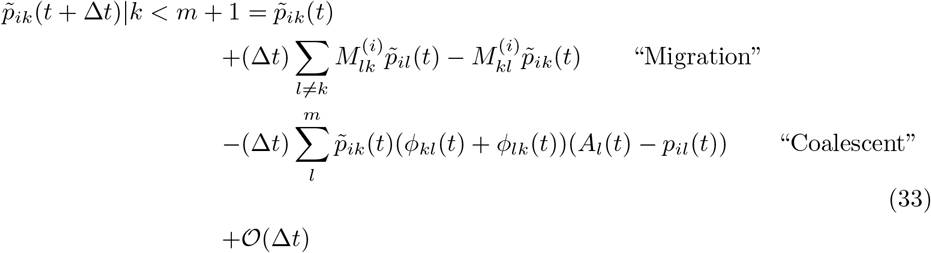

Computing the limit ∆*t* → 0 gives the sytem of differential equations in the main text. Note that the state vector *p*_*j*_ is used instead of 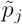 when computing rates of coalescence since we must condition as well on no coalescent event having occured on lineage *j*.

The ‘initial’ state of a lineage following coalescence is found according to the model in [5]. If *i* and *j* coalesce at a lineage *α*, *p*_αk_(*t*) is the probability that *i* was in state *k* and gave birth to *l* or vice versa:

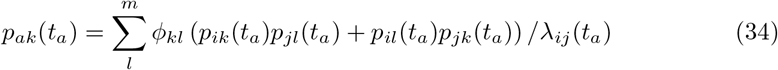

---

**S1 Fig.**
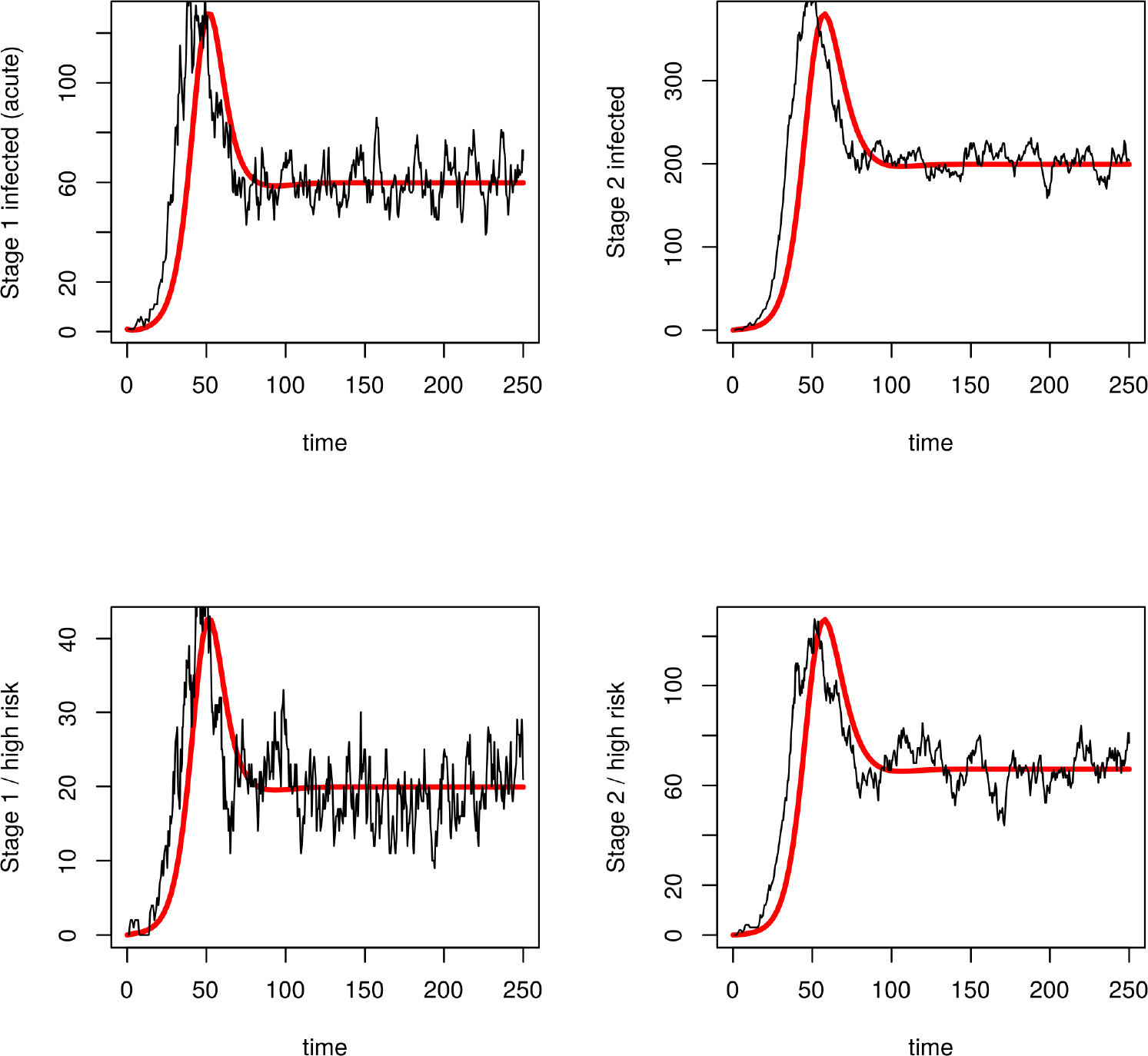
Comparison of stochastic and deterministic trajectories. The stochastic epidemic simulation is shown in black and the deterministic ODE model is shown in red.

## Acknowledgments

The authors thank Tim Vaughan for helpful comments and suggestions. The first version of PhyDyn extended classes from the MASCOT package provided by Nicola Muller.

## References

1. Volz EM, Koelle K, Bedford T. Viral phylodynamics. PLoS Comput Biol. 2013;9(3):e1002947.

2. Drummond AJ, Rambaut A, Shapiro B, Pybus OG. Bayesian coalescent inference of past population dynamics from molecular sequences. Mol Biol Evol. 2005;22(5):1185–1192.

3. Stadler T, Kühnert D, Bonhoeffer S, Drummond AJ. Birth–death skyline plot reveals temporal changes of epidemic spread in HIV and hepatitis C virus (HCV). Proceedings of the National Academy of Sciences. 2013;110(1):228–233.

4. Volz EM, Kosakovsky Pond SL, Ward MJ, Leigh Brown AJ, Frost SDW. Phylodynamics of infectious disease epidemics. Genetics. 2009;183(4):1421–1430.

5. Volz EM. Complex population dynamics and the coalescent under neutrality. Genetics. 2012;190(1):187–201.

6. Frost SDW, Volz EM. Viral phylodynamics and the search for an ‘effective number of infections’. Philos Trans R Soc Lond B Biol Sci. 2010;365(1548):1879–1890.

7. Dearlove B, Wilson DJ. Coalescent inference for infectious disease: meta-analysis of hepatitis C. Philos Trans R Soc Lond B Biol Sci. 2013;368(1614):20120314.

8. Smith RA, Ionides EL, King AA. Infectious Disease Dynamics Inferred from Genetic Data via Sequential Monte Carlo. Mol Biol Evol. 2017;34(8):2065–2084.

9. Anderson RM, May RM, Anderson B. Infectious diseases of humans: dynamics and control. 1992;.

10. Vaughan TG, Kühnert D, Popinga A, Welch D, Drummond AJ. Efficient Bayesian inference under the structured coalescent. Bioinformatics. 2014;30(16):2272–2279.

11. Mueller NF, Rasmussen DA, Stadler T. MASCOT: Parameter and state inference under the marginal structured coalescent approximation; 2017.

12. Beerli P, Felsenstein J. Maximum-likelihood estimation of migration rates and effective population numbers in two populations using a coalescent approach. Genetics. 1999;152(2):763–773.

13. Kühnert D, Stadler T, Vaughan TG, Drummond AJ. Simultaneous reconstruction of evolutionary history and epidemiological dynamics from viral sequences with the birth–death SIR model. J R Soc Interface. 2014;11(94):20131106.

14. Drummond AJ, Bouckaert RR. Bayesian Evolutionary Analysis with BEAST. Cambridge University Press; 2015.

15. Vaughan TG, Leventhal GE, Rasmussen DA, Drummond AJ, Welch D, Stadler T. Directly Estimating Epidemic Curves From Genomic Data; 2017.

16. Lemey P, Rambaut A, Drummond AJ, Suchard MA. Bayesian phylogeography finds its roots. PLoS Comput Biol. 2009;5(9):e1000520.

17. De Maio N, Wu CH, O’Reilly KM, Wilson D. New Routes to Phylogeography: A Bayesian Structured Coalescent Approximation. PLoS Genet. 2015;11(8):e1005421.

18. Rasmussen DA, Boni MF, Koelle K. Reconciling phylodynamics with epidemiology: the case of dengue virus in southern Vietnam. Mol Biol Evol. 2014;31(2):258–271.

19. Volz EM, Ionides E, Romero-Severson EO, Brandt MG, Mokotoff E, Koopman JS. HIV-1 transmission during early infection in men who have sex with men: a phylodynamic analysis. PLoS Med. 2013;10(12):e1001568; discussion e1001568.

20. Volz EM, Ndembi N, Nowak R, Kijak GH, Idoko J, Dakum P, et al. Phylodynamic analysis to inform prevention efforts in mixed HIV epidemics. Virus Evol. 2017;3(2):vex014.

21. Volz E, Pond S. Phylodynamic analysis of ebola virus in the 2014 sierra leone epidemic. PLoS Curr. 2014;6.

22. Ratmann O, Hodcroft EB, Pickles M, Cori A, Hall M, Lycett S, et al. Phylogenetic Tools for Generalized HIV-1 Epidemics: Findings from the PANGEA-HIV Methods Comparison. Mol Biol Evol. 2017;34(1):185–203.

23. Poon AFY. Phylodynamic Inference with Kernel ABC and Its Application to HIV Epidemiology. Mol Biol Evol. 2015;32(9):2483–2495.

24. Karcher MD, Palacios JA, Bedford T, Suchard MA, Minin VN. Quantifying and mitigating the effect of preferential sampling on phylodynamic inference. PLoS Comput Biol. 2016;12(3):e1004789.

25. Rambaut A, Pybus OG, Nelson MI, Viboud C, Taubenberger JK, Holmes EC. The genomic and epidemiological dynamics of human influenza A virus. Nature. 2008;453(7195):615–619.

26. Cori A, Valleron AJ, Carrat F, Scalia Tomba G, Thomas G, Boëlle PY. Estimating influenza latency and infectious period durations using viral excretion data. Epidemics. 2012;4(3):132–138.

27. Dudas G, Carvalho LM, Bedford T, Tatem AJ, Baele G, Faria NR, et al. Virus genomes reveal factors that spread and sustained the Ebola epidemic. Nature. 2017;544(7650):309–315.

28. Koelle K, Rasmussen DA. Rates of coalescence for common epidemiological models at equilibrium. J R Soc Interface. 2012;9(70):997–1007.

29. Rasmussen DA, Volz EM, Koelle K. Phylodynamic inference for structured epidemiological models. PLoS Comput Biol. 2014;10(4):e1003570.

30. Müller NF, Rasmussen DA, Stadler T. The Structured Coalescent and Its Approximations. Molecular biology and evolution. 2017;34(11):2970–2981.

31. Volz EM. rcolgem: statistical inference and modeling of genealogies generated by epidemic and ecological processes. R package version 0.0. 5/r154. 2016;.

32. Biggerstaff M, Cauchemez S, Reed C, Gambhir M, Finelli L. Estimates of the reproduction number for seasonal, pandemic, and zoonotic influenza: a systematic review of the literature. BMC Infect Dis. 2014;14:480.

33. Bettencourt LMA, Ribeiro RM. Real time bayesian estimation of the epidemic potential of emerging infectious diseases. PLoS One. 2008;3(5):e2185.

34. Volz EM, Frost SDW. Scalable relaxed clock phylogenetic dating. Virus Evol. 2017;3(2).

35. Baele G, Li WLS, Drummond AJ, Suchard MA, Lemey P. Accurate model selection of relaxed molecular clocks in Bayesian phylogenetics. Molecular biology and evolution. 2012;30(2):239–243.

36. Palacios JA, Minin VN. Gaussian Process-Based Bayesian Nonparametric Inference of Population Size Trajectories from Gene Genealogies. Biometrics. 2013;.

37. Lemey P, Rambaut A, Welch JJ, Suchard MA. Phylogeography takes a relaxed random walk in continuous space and time. Mol Biol Evol. 2010;27(8):1877–1885.

38. Wakeley J. Coalescent theory: an introduction. 575: 519.2 WAK; 2009.

39. Volz EM, Frost SD. Sampling through time and phylodynamic inference with coalescent and birth–death models. Journal of The Royal Society Interface. 2014;11(101):20140945.

